# Endoplasmic Reticulum contact sites facilitate the coordinated division of *Salmonella-*containing vacuole

**DOI:** 10.1101/2024.05.02.592158

**Authors:** Umesh Chopra, Priyanka Bhansali, Subba Rao Gangi Setty, Dipshikha Chakravortty

## Abstract

*Salmonella* Typhimurium (STM) resides in a membrane-bound compartment called *Salmonella* containing vacuole (SCV) in several infected cell types. Within host cells, the division of bacteria and SCV are synchronous to maintain the single bacterium per vacuole. However, the mechanism regulating the synchronous fission and the machinery is not well understood. The fission of several intracellular organelles is regulated by the dynamic nature of the tubular endoplasmic reticulum (ER). In this study, we have evaluated the role of ER in controlling SCV fission. Interestingly, *Salmonella*-infected cells show the activation of unfolded protein response (UPR) with expanded ER tubules compared to the uninfected cells. Further, changing the expression of ER morphology regulators, such as reticulon-4a (Rtn4a) and CLIMP63, affected bacterial proliferation significantly, suggesting a potential role for tubular ER in facilitating the SCV division. Live-cell imaging analysis shows the marking of tubular ER precisely at the center of the majority of SCV division (78%) sites. We have investigated the role of SteA (a known *Salmonella* effector in modulating the membrane dynamics) in coordinating the SCV division. We observed that SteA resides on the SCV membranes and helps in making membrane contact sites between SCV and ER. Accordingly, the colocalization of ER with SCV enclosing SteA mutant *Salmonella* was significantly reduced compared to SCV-formed by wild-type *Salmonella*. Depletion of *steA* in *Salmonella* resulted in profound defects in SCV division, resulting in multiple bacteria residing in a single vacuole with defects in proliferation compared to the wild-type strain in epithelial cells. Also, during *in vivo* infection, the STM*ΔsteA* mutant shows a defect in colonization in the spleen and liver and affects the initial survival rate of mice. Overall, this study suggests a coordinated role of bacterial effector SteA in promoting the ER contact sites with SCVs and thus regulating the successful division of SCV.

Graphical Abstract

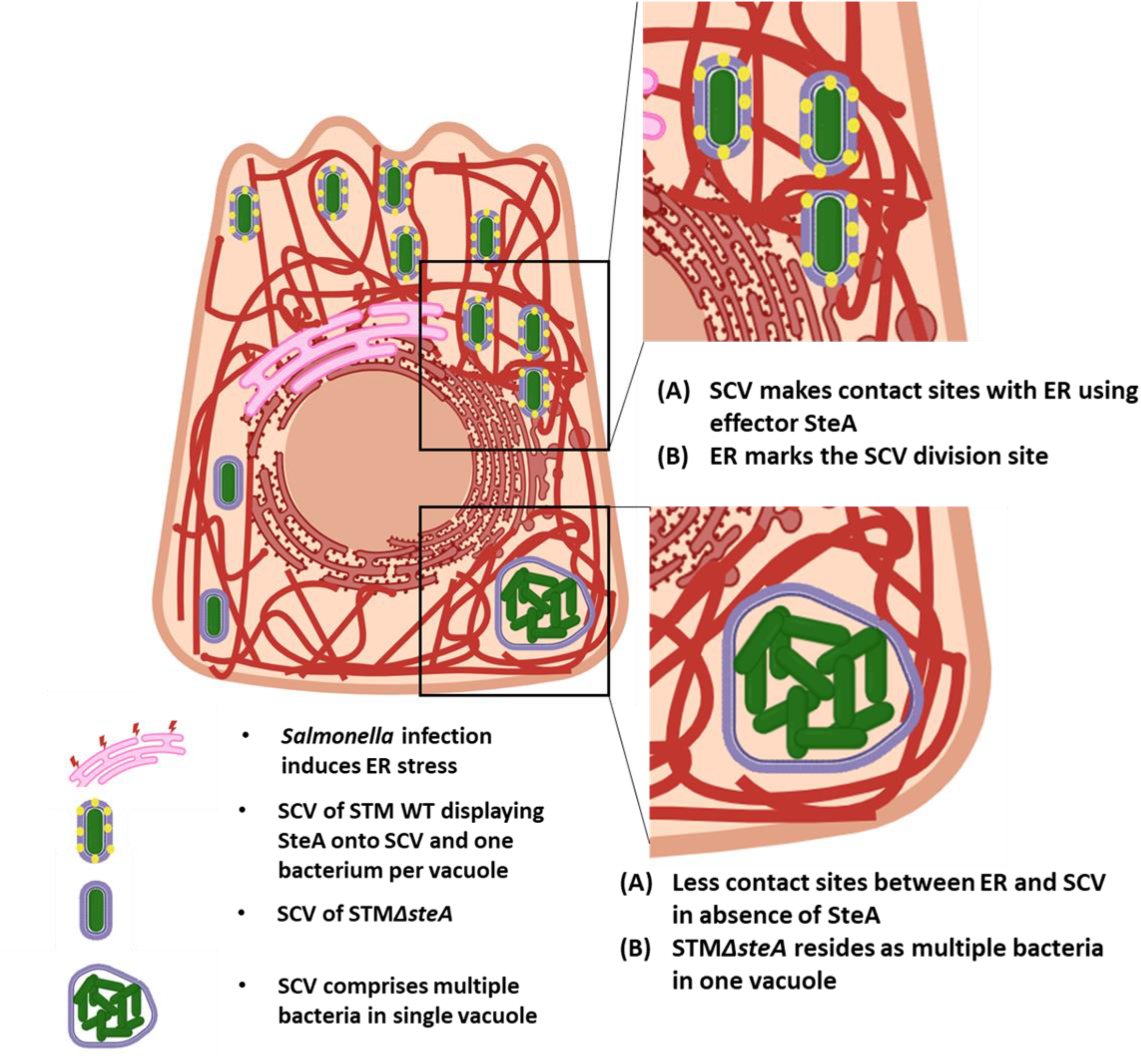

## Introduction

*Salmonella enterica* subsp. *enterica* serovar Typhimurium (STM) is an intracellular pathogen known to infect a wide range of hosts and result in diseases across species, including gastroenteritis to systemic infections [1], [2]. *S.* Typhimurium enters the body through the fecal-oral route following the bacteria utilize an arsenal of strategies to overcome several stresses like the acidic pH of the stomach, gut flora, bile salts, etc., and finally reach the small intestinal epithelia [3]. *Salmonella* secretes several virulence proteins inside the cell, which leads to actin rearrangement and membrane ruffling, which helps in the invasion of bacteria inside the host epithelial cell [4], [5], [6], [7]. *Salmonella* effector SopD is known to facilitate the recruitment of dynamin2 (Dyn2) at the invasion site, which helps in the endocytosis of bacteria inside host cells [8].

Upon invasion into the host cell, *Salmonella* resides in *Salmonella* containing vacuole (SCV), a membrane-enclosed compartment. Mature SCV shares several characteristic markers with lysosomes/late endosomes such as LAMP1 and Rab7 but is negative for acid hydrolases and cathepsins [9]. Inside the epithelial cells, several protruding structures from SCV known as *Salmonella*-induced filaments (SIFs) are observed, which are LAMP1 positive [10], [11]. The membrane composition of SCV is not very well understood, but it contains molecules and proteins from several host organelles and is enriched with several bacterial effector proteins [12], [13]. Our group, as well as several others, have previously shown that *Salmonella* resides as a single bacterium per vacuole state inside the host, suggesting that SCV membrane divides synchronously with the division of *Salmonella* but the mechanism behind SCV division is not known [14], [15], [16], [17]. For successful SCV division, expansion of SCV membrane, pinching of SCV membrane precisely with bacterial division, and lastly, scission of host-derived SCV membrane must be followed in an orchestrated manner.

Several literatures have highlighted the importance of contact sites between cellular organelles, such as contact sites of ER with mitochondria, plasma membrane and endosomes, and contact sites between mitochondria with lysosomes [18], [19], [20], [21]. Contact sites between ER-mitochondria and ER endosomes are very widely studied in the aspect of organelle division [18], [22]. In the case of mitochondria, which comprises an inner and an outer membrane, both membranes divide simultaneously during its fission. The inner mitochondrial membrane uses its machinery for fission, while outer membrane constriction is initiated by the ER, followed by the recruitment of Drp1, which mediates the fission of mitochondria [18], [23], [24]. Bacterial pathogens like *Chlamydia* and *Brucella* make contact sites with cellular organelles to facilitate their proliferation inside the host cell [25], [26], [27]. Interestingly, the SCV membrane also comprises several ER proteins, and recently, membrane contact sites between SCV and ER have also been observed [28]. To understand the mechanism behind the SCV division, we reasoned that SCV is partially like mitochondria. SCV and mitochondria (being an endosymbiont) encompass an outer membrane derived from the eukaryotic host, while the inner membrane of mitochondria is prokaryotic in nature, like the cell wall of *Salmonella*. To understand the same, we hypothesized that a process similar to mitochondria could lead to SCV division. Our results suggest that *Salmonella* infection leads to activation of UPR, ER expansion, and an increase in ER tubulation. We have observed ER tubules mark at the fission site of SCV division which possibly initiate the constriction. *Salmonella* translocated effector SteA localizes onto the SCV membrane and possibly helps in making contact with ER, suggesting that SteA can act as a factor to recruit the ER onto SCV and thus facilitate the SCV division. STM*ΔsteA* resides as multiple bacteria in one big vacuole with defects in proliferation. During *in vivo* infection, STM*ΔsteA* shows defects in colonization in the spleen and liver if injected through the intraperitoneal route. Also, there is a delay in the onset of mice death upon infection with STM*ΔsteA* as compared to STM WT, but eventually, mice succumbed to death on the 8th day post-infection. Overall, our study unravels, for the first time, the coordinated role of ER and bacterial effector SteA in successful SCV division inside the cell.

## Materials and Methods

### Bacterial strains and culture conditions

*Salmonella enterica* subspecies *enterica* serovar Typhimurium (STM WT) wild-type strain ATCC 14028s was used in all experiments. The bacterial strain was cultured in Luria broth (LB-Himedia) with constant shaking (170 rpm) at 37°C orbital-shaker. Antibiotics like kanamycin (50 µg/ml) and ampicillin (50µg/ml) were used wherever required. A one-step gene inactivation method was used for the generation of knockout strains [29]. Wild type as well mutant strains were transformed with pFPV-m-cherry or pFPV-GFP plasmid for immunofluorescence assays.

#### Generation of Complement strain and SteA-eGFP plasmid

Low copy number plasmid pQE-60 was used to prepare the SteA-HA complement strain. Briefly, PCR amplification (using Thermo Q5 DNA polymerase) of the *steA* gene was performed using Wild Type STM colony with respective PCR primers. Forward cloning primer was chosen from a position of 250 base pairs upstream of the gene start site to amplify the native promoter of the gene during PCR and to maintain the native expression level of protein, and the sequence of HA tag was added in reverse primer. The amplified PCR product was purified using a Gel extraction kit (Qiagen). The PCR product and Plasmid were subjected to restriction digestion by using Xho1 (NEB) and Hind III (NEB) at 37°C for 2 hours. The double-digested PCR product and plasmid were further subjected to Ligation using T4 DNA ligase (NEB) at 16°C for 12-14 hours. The ligated product was further electroporated in STMΔ*steA* strain to prepare the complemented strain, i.e., STM*ΔsteA*:pQE60-*steA* (hence referred to as STM*ΔsteA:steA*). The expression level of complement was confirmed with western blotting and confirmed with DNA sequencing.

For preparation of SteA-eGFP plasmid for transfection study, eGFP-N3 empty vector was used. The sequence of *steA* was amplified via PCR from STM WT using primer with restriction site for Xho1 and BamH1. The entire process for cloning was performed as discussed above.

**Table 1.**
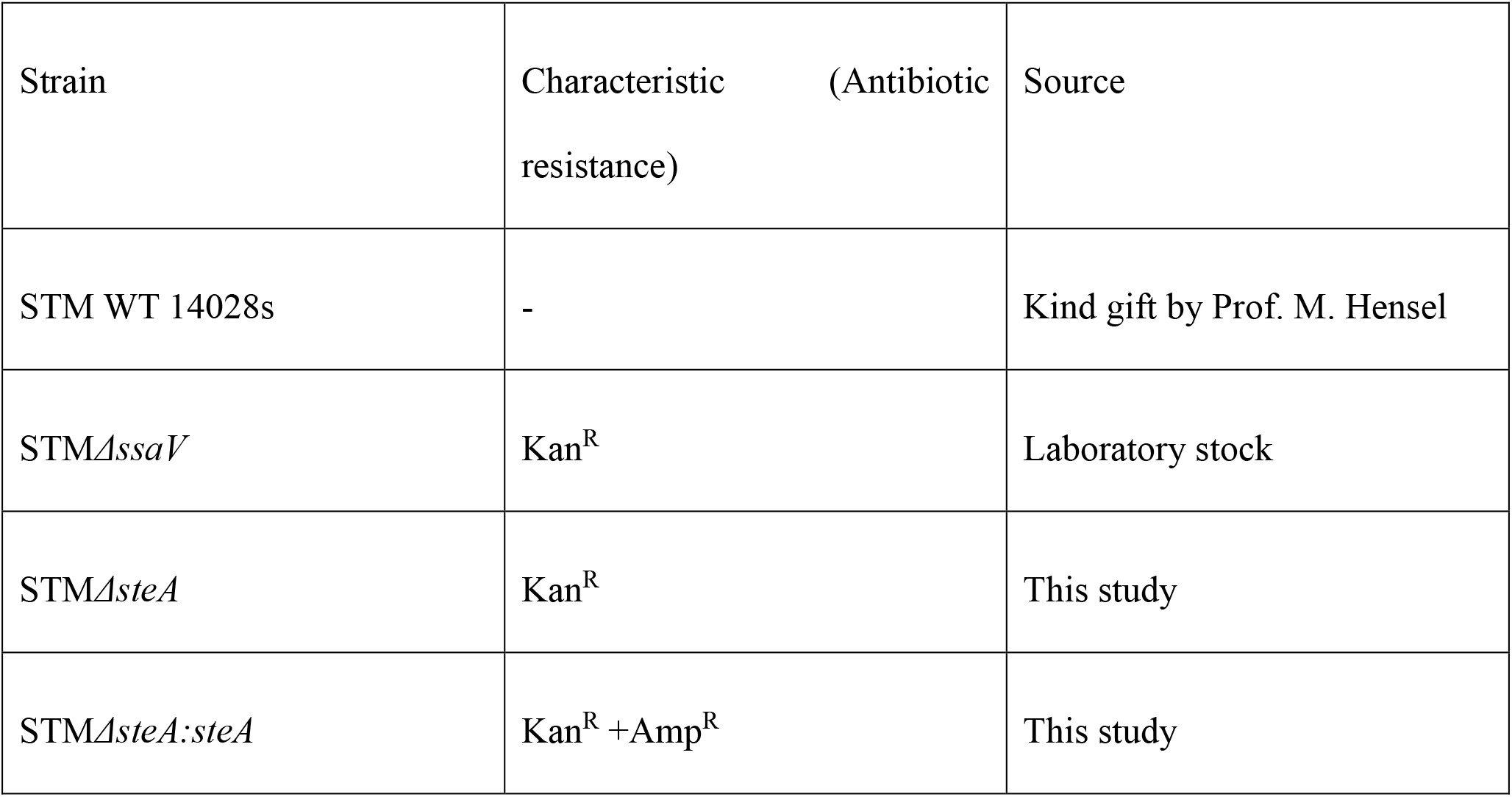
List of bacterial strains used in this study

| Strain | Characteristic (Antibiotic resistance) | Source |
| --- | --- | --- |
| STM WT 14028s | - | Kind gift by Prof. M. Hensel |
| STMΔ*ssaV* | Kan<sup>R</sup> | Laboratory stock |
| STMΔ*steA* | Kan<sup>R</sup> | This study |
| STMΔ*steA:steA* | Kan<sup>R</sup> +Amp<sup>R</sup> | This study |

#### Eukaryotic Cell Culture, Transfection and Plasmids

HeLa cells and RAW 264.7 murine macrophages were cultured in DMEM (Lonza) containing 10% FBS (Gibco) at a humified incubator, maintaining a temperature of 37°C and 5% CO_2_. Cells were seeded in 6 wells, 12 wells, or 24 well plates at a confluency of 60-70% prior to infection. Cells were maintained in the presence of 1% penicillin and streptomycin (penstrep). Cells were given a PBS wash and supplemented with fresh media without penstrep at least 6 hours prior to infection. Plasmid overexpression or knockdown via shRNA was done using Lipofectamine 3000 using manufacturer protocol. Briefly, 300ng of plasmid was mixed in opti-MEM media for 5 min while 1ul of L3000 was added (for one well of 24 well plates) in opti-MEM for 5 min separately and further mixed and incubated for 20 min at room temperature. Post incubation, this mixture was added to cells for 6 hours for transfection to take place. Cells were washed with PBS and supplemented with complete media post 6 hours of treatment, and post 48h of transfection, infection was given. Overexpression constructs such as pHAGE2 mCherry-Rtn4a was a kind gift from Prof. Tom Rapoport (Addgene plasmid 86683) [30] mCh Climp-63 and mCh sec61β was a kind gift from Prof. Gia Voeltz (Addgene plasmid 136293, Addgene 49155) [31], [32], RFP KDEL construct was a kind gift from Prof. Nagaraj Balasubramanian [33], GFP-LAMP1 was provided by Prof. Mahak Sharma [34]. The shRNA plasmids were obtained from the human genome-wide TRC shRNA library (purchased from Sigma-Aldrich, USA, Catalog no. SH0411), available at shRNA resource centre, MCB, IISc. pLKO.1-puro non-mammalian shRNA (SHC002, shControl) was used as a control in all the shRNA mediated knockdown experiments. Plasmid sequences and details are added in supplementary table 2.

#### Protocol for infection (Gentamicin Protection Assay)

The cells were seeded at the required confluency and infected with *Salmonella* strains with actively growing log phase culture (for epithelial cells) and stationary phase culture (for RAW 264.7 macrophages) at Multiplicity of infection (MOI) 10 for Intra Cellular Survival Assay (ICSA) and confocal imaging. For the live cell imaging experiment, an MOI of 25 was used to increase the event of finding infected cells for imaging. For confocal microscopy studies, cells were seeded on sterile glass coverslips at least 24 hours before infection. Upon infecting the cells at the required MOI, the plate was centrifuged at 600 rpm for 5 min to facilitate the adhesion, and then the plate was incubated for 25 min at 37°C and 5% CO_2_. Post incubation, bacteria-containing media was removed, and cells were given 2 PBS washes to remove any extracellular bacteria. Fresh media containing 100µg/ml of gentamicin was added into wells and incubated at 37°C for one hour. Following this, cells were again given 2 PBS washes, and fresh media containing 25µg/ml of gentamicin was added, and this concentration was maintained till the end point of the experiment. Cells were harvested at required time points, such as 2h and 16h for ICSA, 2h, 6h, and 16h for confocal microscopy-based experiments.

#### Confocal Microscopy

After an appropriate hour post-infection, cells were washed with PBS twice and fixed with 3.5% PFA at room temperature for 15 min. After fixation, coverslips were washed with PBS, and cells were stained with primary antibody prepared in 2% BSA in PBS carrying 0.01% of saponin (for membrane permeabilization) for 1 hour at room temperature or 4°C overnight in a cold room as required. Post incubation, coverslips were washed again twice with PBS and incubated with specific secondary fluorophore-labeled antibodies and incubated for one hour at room temperature. Coverslips were then mounted on clean glass coverslips using mounting media. Once the mounting media was dried, coverslips were stabilized by sealing the periphery with transparent nail paint on the sides. Images were acquired on a confocal scanning laser microscope (Zeiss 880 multiphoton microscope or Leica sp8 falcon system with or without zoom as required). Zen black software was used for all colocalization-based quantification and to calculate mean fluorescence intensity (MFI).

Calculations of CTCF, percent of ER tubules, SCV diameter, and Line scans were prepared using ImageJ. The ER tubule percentage is calculated by calculating the sheet percentage from the total cell area. Sheet area / total area *100 = Sheet area percentage and further 100 - Sheet area percentage = tubule area percentage [35]. Live cell microscopy was performed on the Leica SP8 Falcon microscope, and live cell video was processed and prepared using LAS X software. Antibodies used in this study are mentioned in supplementary table 3.

#### Bacterial enumeration after infection for Intracellular survival assay

After the appropriate hour of post-infection, cells were washed with PBS at RT and lysed using 0.1% Triton-X 100. Cell lysate was plated on SS agar to obtain colony forming unit (CFU) and it was used for the calculation of Percent Invasion and Fold proliferation using this formula.

Percent Invasion: (CFU obtained at 2 hours post-infection/ CFU obtained from pre-inoculum) *100 Fold proliferation: CFU obtained at 16-hour post-infection/CFU obtained at 2 hours PI

#### RNA isolation and Real-Time PCR (qRT-PCR)

RNA isolation was performed using TRIzol (Takara) following the manufacturer’s protocol. Quantification of RNA was done using nano-drop (Thermo-Fisher Scientific) and RNA samples were run on 1% agarose gel to check the quality of RNA. 2 µg of RNA was subjected to DNase I (Thermo Fischer Scientific) treatment at 37°C for 1 hour followed by addition of 0.5M EDTA (final concentration 5mM) and heat inactivation at 65°C for 10 mins to inactivate the enzyme. Further, cDNA was prepared by using a cDNA synthesis kit (Takara) as per the manufacturer’s instructions. The expression profile of all genes of interest was evaluated using RT primers by using SYBR green mix (Takara) in a Biorad Real-Time PCR machine. The expression value of the target gene was normalized to housekeeping internal control β-actin and further compared with uninfected cells. Primers used in this study are mentioned in Supplementary Table 4.

### *In vivo* animal experiment

6-8 weeks old C57BL/6 female mice were infected by orally gavaging 10^7^ CFU of STM WT, STM*ΔsteA* and STM*ΔsteA:steA*. 5 days post-infection, mice were euthanized, and organs such as liver, spleen, MLN, and blood were harvested to study the colonization in organs. For intraperitoneal injection, 10^3^ CFU of bacteria was administered per animal, and 3 days post-infection, organs were homogenized and further plated on *Salmonella*-*Shigella* agar. Obtained CFU values were normalized with the gram weight of the organ and then converted into a log scale. Blood was collected through heart puncture and plated on SS agar. For the mice survival assay, 10^8^ CFU was administered to each mouse via oral gavage. Mice were monitored every day till the time none of the mice in the cohort survived. The animal experiments were carried out in accordance with the approved guidelines of the institutional animal ethics committee at the Indian Institute of Science, Bangalore, India (Registration No: 48/1999/CPCSEA). All procedures involving the use of animals were performed according to the Institutional Animal Ethics Committee (IAEC)-approved protocol.

#### Statistical analysis

Statistical analysis was performed using GraphPad Prism software. For analysis, the student’s t-test (unpaired parametric test) was performed. The results are indicative of mean± SD or mean± SEM. To calculate the statistical significance of bacterial colonization *in vivo*, the Mann-Whitney test was performed. Group size, experimental number, no of cells in each set, and P value of each experiment are described in figure legends.

## Results

### *Salmonella* infection leads to the activation of UPR and expansion of the Endoplasmic Reticulum

Several bacterial and viral infections are known to activate UPR for their survival [36], [37]. UPR helps to alleviate the ER stress by initiating several transcriptional and translational regulations which lead to the maintenance of ER homeostasis [38], [39]. Inside the cell, UPR signaling is maintained by transmembrane proteins located in the ER membrane and their catalytic domain towards the cytosol. Inositol requiring enzyme-1 (IRE-1), Protein kinase c like ER kinase (PERK), and Activating transcription factor-6 (ATF-6) comprises the main arm of UPR. Recent reports have shown the contact of SCV with ER, also the presence of ER proteins in the membrane fraction of SCV suggests the crosstalk between them [12], [28]. Here, we investigated how *Salmonella* infection modulates the ER dynamics. To understand the activation of UPR by any of these pathways, we used confocal microscopy and quantified the translocation of downstream transcription factors, such as XBP1, ATF4, and ATF6, inside the nucleus upon infection. SPI2 mutants of *Salmonella*, STM*ΔssaV* (which is unable to secret SPI-2 encoded effector protein in host cytosol) was kept as a control to understand if activation of UPR is mediated by bacterial effectors or dependent on bacterial invasion. Our results suggest that from as early as post 2h of *Salmonella* infection, both the transcription factors, ATF4 and ATF6 from the cytosol translocated to the nucleus of infected cells as compared to uninfected cells and STM*ΔssaV* infected cells, suggesting that only wild-type *Salmonella* can activate UPR (fig 1A-1D). *Salmonella-*induced ATF4 and ATF6 translocation were retained till 16h post-infection (fig S1A-S1D). However, we didn’t observe any increase in XBP1 recruitment inside the nucleus (fig 1E,1F) upon infection at any time point post-infection (fig S1E, S1F), suggesting that *Salmonella* infection upregulates UPR by PERK and ATF6 pathways in HeLa cells. To alleviate the ER stress, activation of UPR leads to the expansion of the ER, protein folding capacity, and increased phospholipid production [40]. Increased bacterial proliferation requires a continuous source of membrane for the increasing number of SCV, and increased phospholipid production upon UPR could facilitate the SCV proliferation inside the host cell. Therefore, we sought to determine if *Salmonella* infection led to ER expansion during the pathogenesis phase. Our transcriptional analysis suggests that upon infection, there is upregulation at the transcript level of ER shaping proteins such as *Climp63* (responsible for maintaining ER sheets) and *Reticulon-4a* (*Rtn4a*, essential for maintaining ER tubules) (fig 1G, fig 1H). The upregulation was significantly observed at the late time point of infection, i.e., 16h post-infection, suggesting that active proliferation of *Salmonella* and not just invasion (as evident at 2h and 6h post-infection) leads to ER expansion.

**Fig. 1:**
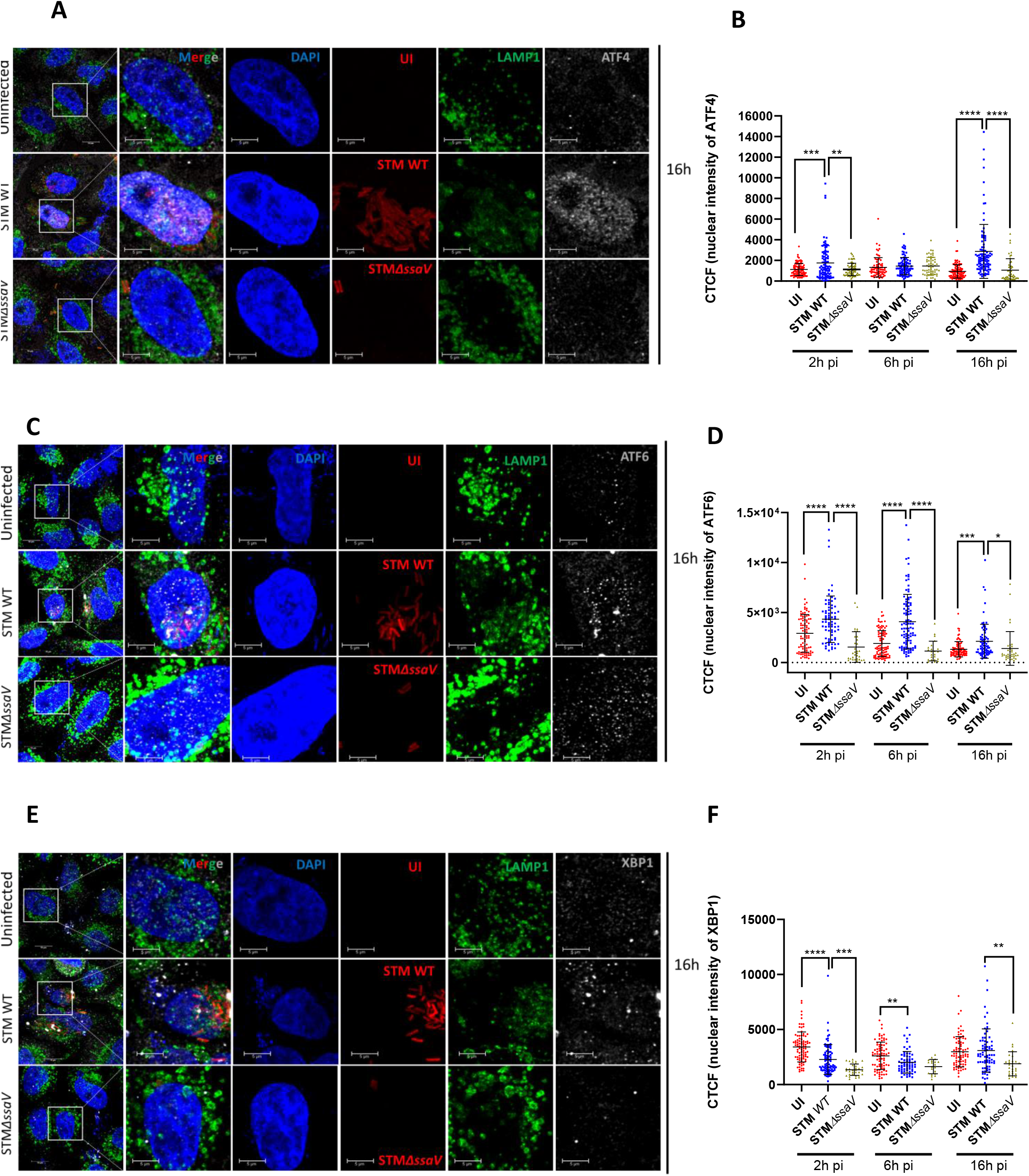
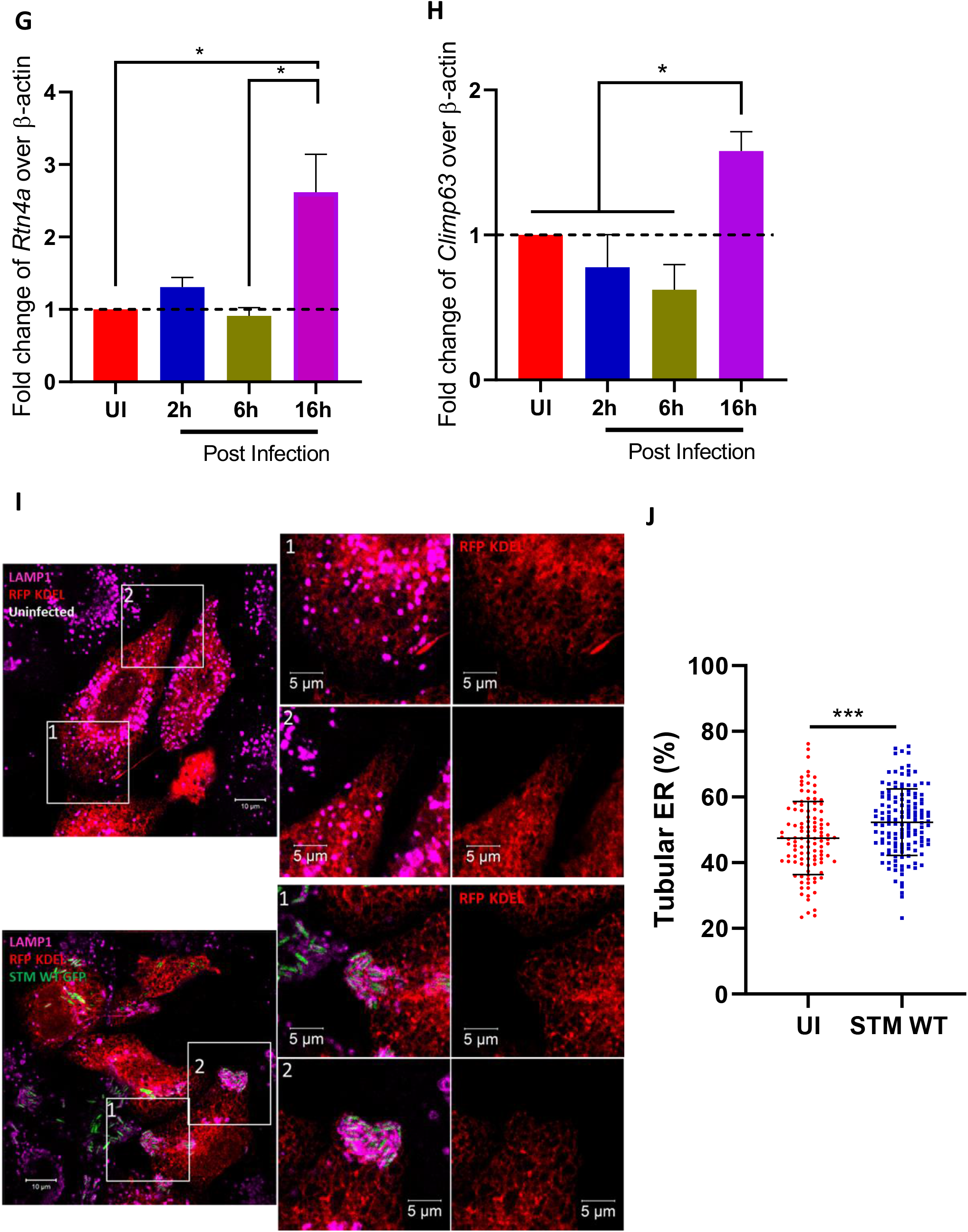
**Salmonella infection leads to the activation of UPR and expansion of the Endoplasmic Reticulum** (A) Representative confocal microscopic images of uninfected and infected HeLa cells with STM WT mCh or STM*ΔssaV* mCh at MOI 10, 16h post-infection and immuno- stained against ATF4, LAMP1 and DAPI. (B) Represents the quantification of Cell Total Corrected Fluorescence (CTCF) intensity of ATF4 inside the nucleus, data is representative of n=100-120 cells, from 4 independent experiments, mean±SD (C) Representative confocal microscopic images of uninfected and infected HeLa cells with STM WT mCh or STM*ΔssaV* mCh at MOI 10, 16h post infection and immuno-stained against ATF6, LAMP1 and DAPI. (D) Represents the quantification of Cell Total Corrected Fluorescence (CTCF) intensity of ATF6 inside the nucleus, data is representative of n=60-90 cells from 3 independent experiments, mean±SD (E) Representative confocal microscopic images of uninfected and infected HeLa cells with STM WT mCh or STM*ΔssaV* mCh at MOI 10, 16-hour post-infection and immuno- stained against XBP1, LAMP1, and DAPI (F) Represents the quantification of Cell Total Corrected Fluorescence (CTCF) intensity of XBP1 inside the nucleus, data is representative of n=60-90 cells from 3 independent experiments, mean±SD (G) Quantitative real-time PCR of *Rtn4a* (H) and *Climp63*, N=3, n=3, mean±SEM (I) Confocal microscopic images of Un-infected and infected HeLa cell showing ER expansion (J) Represents the quantification of tubular ER, data is representative of n=100- 150 cells from 4 independent experiments, mean±SD. Student’s t-test was used to analyze the data ****<0.0001, ***<0.001, **<0.01, *<0.05.

Since an increase in the transcript levels of ER shaping proteins was observed at the later time of infection, we tried to quantify the percentage of ER sheets and tubules using the ER marker, RFP- KDEL, upon infection to understand ER expansion. Upon analysis of microscopy images, we observed that *Salmonella*-infected cells showed an increased percentage of ER tubules compared to uninfected cells (fig1I, fig1J). The changes in percent ER tubules were only evident in infected cells as compared to uninfected and bystander cells (neighboring cells that are not infected), suggesting that stabilized SCV and bacterial effectors are responsible for this phenotype (fig S1G and S1H). Together, this data indicates that *Salmonella* infection leads to changes in ER morphology.

#### Expansion of ER tubules facilitate SCV proliferation and its division

Dynamic ER tubules play an essential role in forming membrane contact sites with several organelles [41], [42]. Figure 1 indicates that actively proliferating bacteria increased tubular ER and upregulated the expression of *Rtn4a*. We also tried to understand if there is an advantage of the increased tubules to bacterial proliferation. To identify the role of ER tubules in mediating bacterial proliferation, we transiently overexpressed reticulon 4a (Rtn4a) in HeLa cells, which helps to increase ER tubules [43], and then quantified the proliferation of bacteria by intracellular survival assay (ICSA). As compared to empty vector and general ER marker sec61β-mCh control, increased ER tubulation by Rtn4a led to the higher proliferation of *Salmonella* with no significant change in percent invasion, suggesting an increase in ER tubules helps in bacterial proliferation (fig 2A, figS2A). To validate if this change in proliferation was dependent solely on the function of Rtn4a, knockdown of Rtn4a via shRNA was performed (figS2B). Knockdown of Rtn4a (῀40% reduction at transcript level) led to reduced proliferation of bacteria as compared to scrambled control, hence strengthening the crucial role of ER tubules in bacterial proliferation (fig2B). Climp63 overexpression led to an increase in ER sheets as opposed to tubules. We quantified the number of SCV per cell upon transiently overexpressing the Climp63 in HeLa cells. Our data indicated that the number of SCV present per cell decreases upon overexpressing ER sheets using Climp63 compared to controls (fig2C, fig2D). Division of *Salmonella* is linked with the fission of SCV membrane [15]. Since the intricate balance of ER sheets and tubules affects bacterial proliferation, we hypothesized that ER tubules might facilitate SCV fission. In the case of mitochondria, membrane-less organelle, and endosomes, ER marks the division site, and later fission is completed by Drp1 (in mitochondria) and coronin1c and TMCC 1 complex (in endosome) [18], [24], [44], [45]. To investigate our hypothesis of ER tubules playing a role in SCV fission, we performed live cell imaging in cells transfected by RFP-KDEL (to mark the ER) followed by infection with *Salmonella*. Quantification of data suggests that ER tubules mark the division site precisely at the center of dividing SCV in 78% of events, as observed by image snapshots and corresponding line scans (fig 2E-G, video S1). STM WT infected RAW macrophage also showed contact of ER on the dividing bacteria (video S2, S2C). This was also evident in fixed cell images where we could observe Lamp1 enclosed *Salmonella* division in SCV marked by ER tubules exactly in the center in 56.7±6.61% of fission events, however, in 14±9.07% of events, ER was seen in proximity to the center and hence mentioned as adjacent while 29.2±8.4% of dividing SCV did not possess ER tubules on them (fig 2H, fig2I). This data suggests the importance of ER tubules in marking the fission site of SCV. Taken together this data suggests that ER tubules help in the proliferation and division of SCV by marking the fission site inside the cell and help in maintaining SCV number within the cell.

**Fig. 2:**
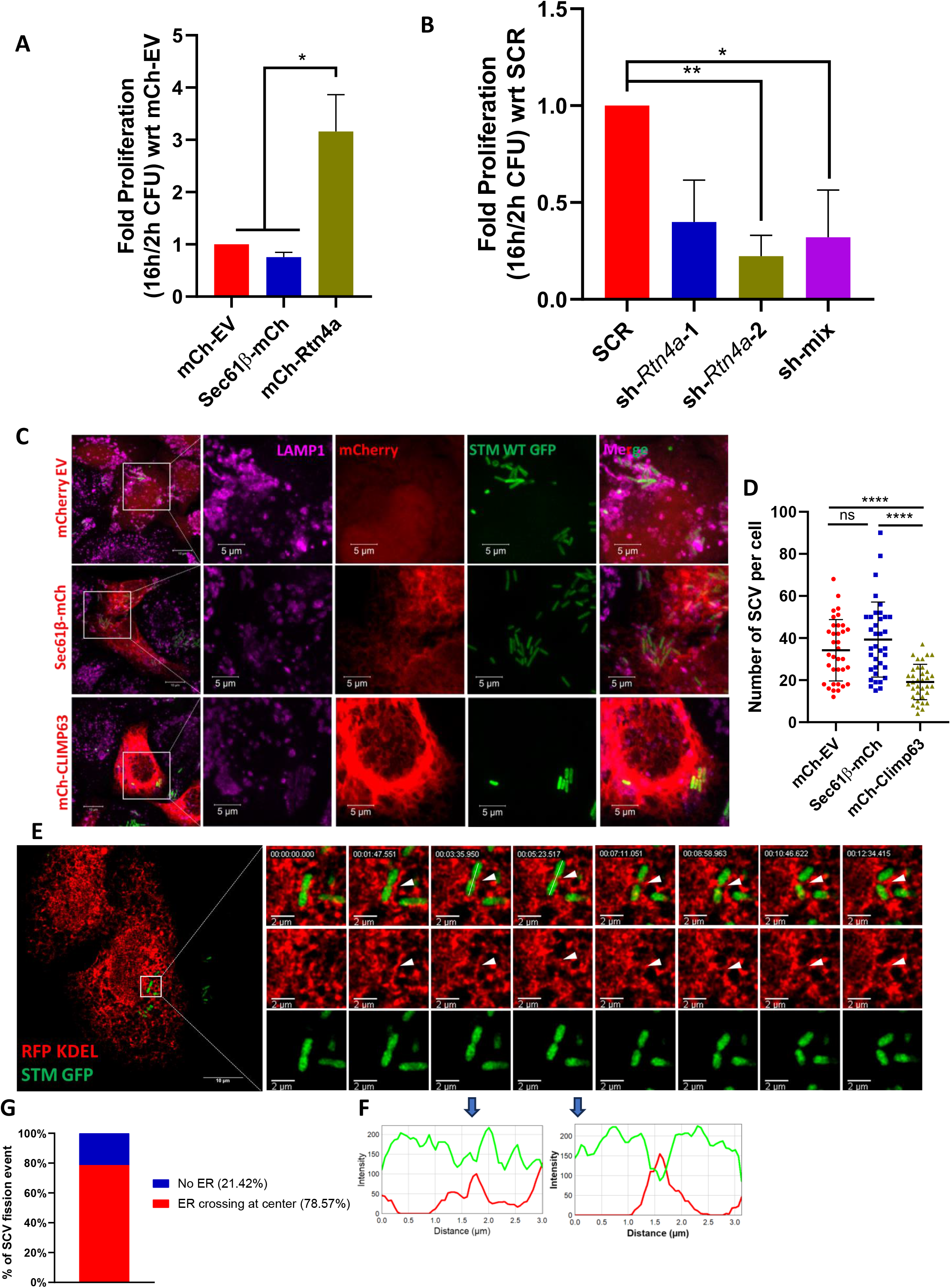
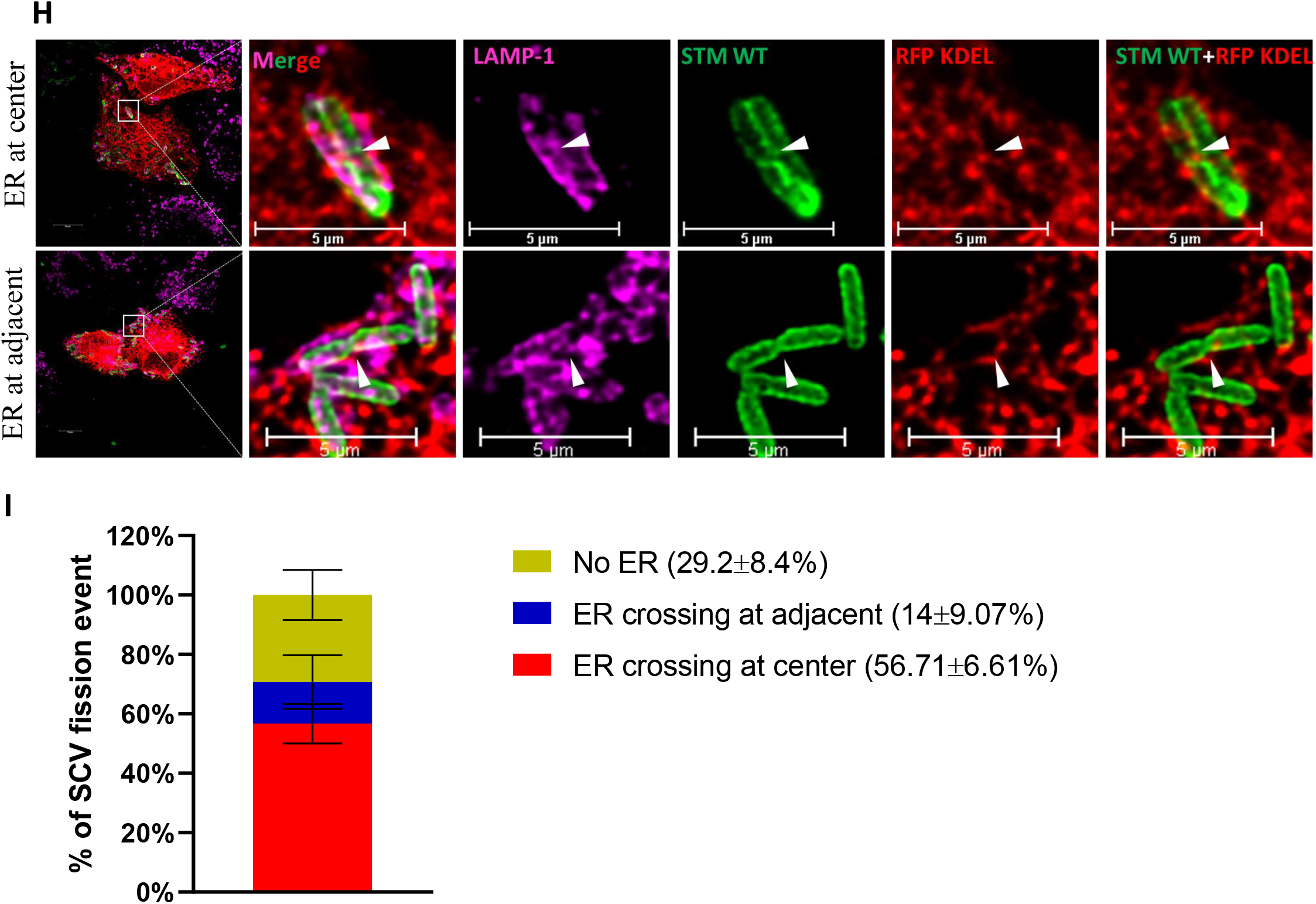
**Expansion of ER tubules facilitates SCV proliferation and its division** (A) Represents the ICSA upon overexpression of Rtn4a, data is representative of N=3, n=3 mean± SEM (B) ICSA upon Rtn4a knockdown in HeLa cells, data is representative of N=3, n=3 mean± SEM (C) Representative confocal microscopy images of SCV number per cell upon overexpression of mCh-EV, sec61β-mCh, and mCh-Climp63, 16h post infection (D) Quantification of the number of SCV per cell, 16h post infection, data is representative of n=40-60 cells from 2 independent experiments (E) Snapshot of live cell imaging of *Salmonella* infected HeLa cells, White arrowhead represent the ER overlap at dividing bacteria. White line along the length of bacteria represent the portion used for line scan. (F) Showing the line scan of images. Blue arrows represent the snapshot for which the line scan is analyzed (snapshots 3 and 4) (G) Quantification of SCV fission events marked by ER, data is representative of n=11 SCV fission events from 4 independent experiments (H) Confocal microscopic images representing SCV fission events marked by ER, at center or adjacent (I) Quantification of fission events in fixed images marked by ER, data is representative of n=49 SCV fission events from 4 independent experiments. White arrowhead represent the ER overlap at dividing SCV. Student’s t-test was used to analyze the data ****<0.0001, ***<0.001, **<0.01, *<0.05.

#### *Salmonella* translocated effector SteA is crucial for maintaining SCV contact site with ER, resulting in a single bacterium per vacuole

SCV division leads to a single bacterium per vacuole throughout the course of pathogenesis in cell lines or *in vivo* [14]. Domingues et al., suggested the role of SteA in maintaining the membrane dynamics of SCV [46]. In accordance with the same, we also observed that *steA* mutant strain of *Salmonella* (STM*ΔsteA*) resides as multiple bacteria in one single vacuole (fig 3A). Our data suggests that compared to STM WT, 30±3% of mutant bacteria reside in one big vacuole, suggesting that loss of function of *steA* leads to improper SCV fission (fig 3B). Live cell imaging shows several points of contact between SCV and ER, suggesting that SCV actively makes membrane contact sites with ER for its survival and proliferation (fig 2E, fig 3C). SteA resides onto the PI4P-enriched SCV membrane (fig 3C), and hence, we asked if membrane contacts between ER and SCV are maintained by SteA. Microscopy images show that SteA-enriched SCV makes contact with ER (fig 3C). Quantification of the data suggests that ER colocalizes more with the SCV of STM WT than the SCV formed by STM*ΔsteA* (as observed colocalization between SCV marker Lamp1 and RFP KDEL), suggesting that SteA helps in making contact sites of SCV with ER (fig 3D, fig 3E). This was also observed upon transfecting HeLa cells with the SteA-eGFP construct, where SteA-eGFP showed higher colocalization with ER marker RFP-KDEL as compared to eGFP empty vector control, strengthening the fact that SteA possibly associate with ER (figS3A, S3B). Overall, this data suggests that SteA might help in making contact sites with ER and further recruiting ER onto the SCV. These contact sites can be used as a stealthy approach by *Salmonella* to recruit membrane from ER, and along with other *Salmonella* effector proteins, facilitating SCV fission.

**Fig. 3:**
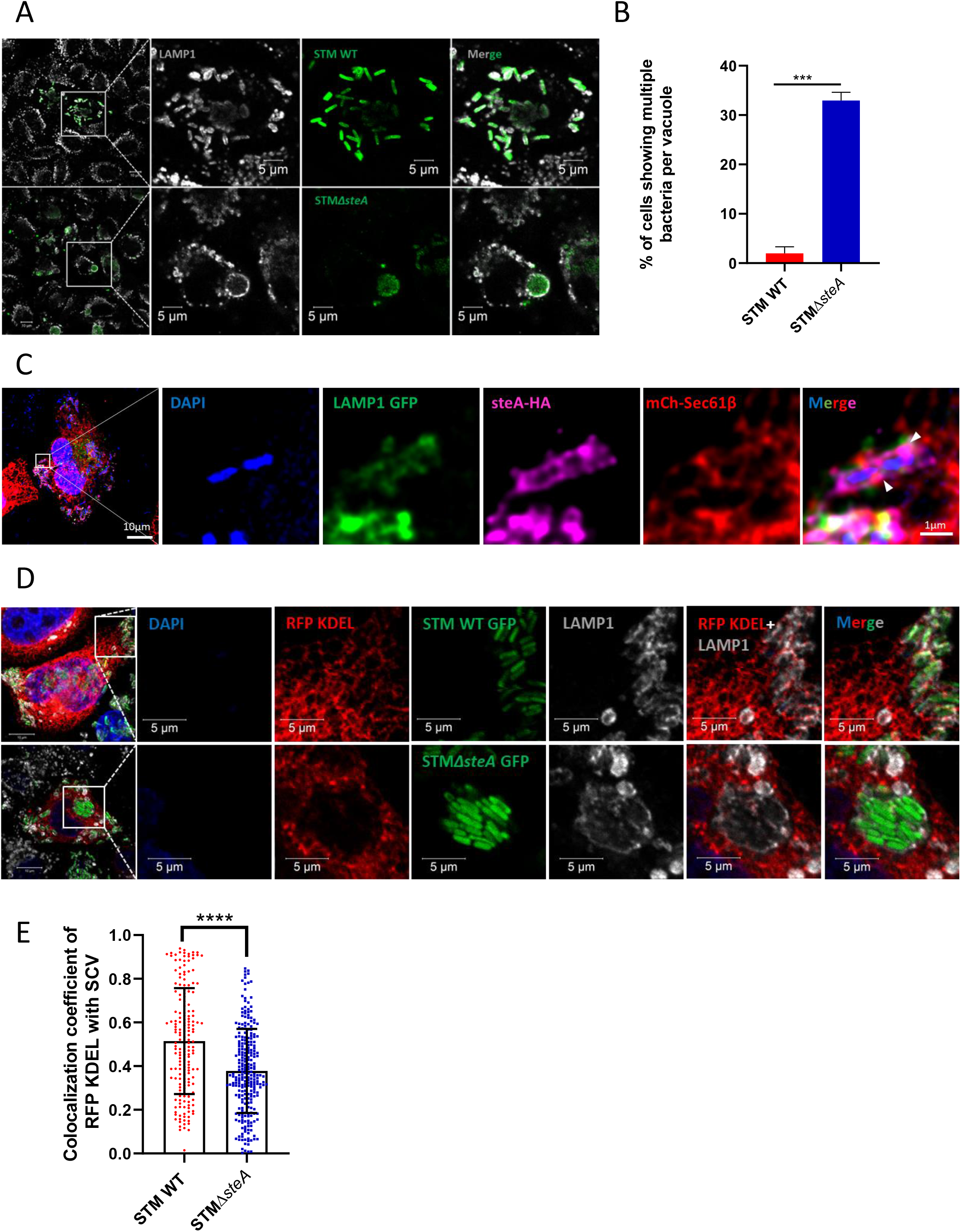
**Salmonella translocated effector SteA is crucial for maintaining SCV contacts with ER, resulting in a single bacterium per vacuole** (A) Represents confocal microscopy images showing SCV formed by STM WT and STM*ΔsteA,* 16h post-infection (B) Quantification of the number of cells showing multiple bacteria in one vacuole, 16h post-infection, data is representative of n=120-150 cells from 3 independent experiments, mean±SEM (C) Representative confocal image of infected HeLa cell showing contact between SteA-HA and ER, arrowheads represent interaction between ER and SteA-HA (D) Confocal images represents the colocalization of ER with SCV formed by either STM WT or STM*ΔsteA,* 16h post-infection (E) Quantification of colocalization coefficient of ER marker RFP KDEL with SCV marker LAMP1, Data is representative of n=80-100 cells from 2 independent experiments, 16h post infection, mean±SD. Student’s t test was used to analyse the data ****<0.0001, ***<0.001, **<0.01, *<0.05.

#### STM*ΔsteA* shows a defect in proliferation and pathogenicity *in vivo*

Previous reports from our lab suggested that *Salmonella* utilizes the strategy of SCV division to increase the number of SCV per cell, so it becomes difficult for the cellular system to eradicate the growing SCV [14]. Since *steA* mutants form a big vacuole inside the cell, we asked how the mutant strain proliferates in the cell. As compared to STM WT and STM*ΔsteA:steA* (complemented strain), STM*ΔsteA* proliferated significantly less in the HeLa cell line. (fig 4A). Less proliferation of mutant strain suggests that those bulky vacuoles might be targeted by cellular machinery for degradation. Quantification of vacuole size suggests that the diameter of SCV formed by STM*ΔsteA* is around 6.9±3.13µm whereas SCV of STM WT is 0.4±0.11µm (fig 4B, fig 4C). Using LC3B as a marker to detect the extent of autophagy, we observed that LC3B colocalized more with SCV of *steA* mutants as compared to SCV of STM WT at 16h post-infection (fig4D, fig4E). Actively proliferating *Salmonella* also reduces the number of lysosomes inside the cell to escape its fusion with acidic lysosomes [14]. In STM*ΔsteA* infected cells, we observed a higher intensity of Lamp1 similar to uninfected cells, suggesting that STM*ΔsteA* is unable to change the lysosomal dynamics as opposed to STM WT. A higher flux of lysosomal proteins, along with the recruitment of LC3B, led to less proliferation of the STM*ΔsteA* inside the cell (figS4A, figS4B). Next, we were interested in understanding the pathogenicity of STM*ΔsteA in vivo*. For this, we intraperitoneally injected 10^3^ CFU of *Salmonella* per C57BL/6 mice, and 3^rd^-day post-infection mice were euthanized, and organs such as liver and spleen were harvested to observe organ burden. Our data suggest that 3rd-day post-infection, STM*ΔsteA* shows less colonization in the spleen and liver as compared to STM WT and STM*ΔsteA:steA* (fig 4G, fig4H). To further understand the pathogenicity of the mutant strain, we orally gavaged C57BL/6 mice with 10^7^ CFU of STM WT, STM*ΔsteA*, and STM*ΔsteA:steA,* and 5 days post-infection, mice were euthanized, and organs such as liver, spleen, and mesenteric lymph nodes were harvested since they act as a secondary site of infection (fig 4J-4L). We observed no significant change in the colonization of bacteria in these organs, suggesting STM*ΔsteA* doesn’t lose its capability to cross the gut barrier and colonize and disseminate in distal organs and blood if injected through the oral route (figS4C). However, upon performing a survival assay of mice upon infection through oral gavage, we noticed the delayed onset of mice death in the STM*ΔsteA* infected cohort (fig4M), where STM WT infected mice started succumbing to infection from 5^th^ day post-infection, STM*ΔsteA* infected cohort started succumbing to infection at 7^th^ day post-infection, and at 8th-day post-infection, all the mice succumbed to death. Taken together, this data suggests that if the fission of SCV is altered, *Salmonella* resides as multiple bacteria in one big vacuole and proliferates less. Those big/bulky vacuoles are further targeted by autophagic machinery and cleared by active lysosomes in the cell culture system. During *in vivo* infection through the intraperitoneal route, STM*ΔsteA* shows defects in colonization in the spleen and liver and also affects the initial survival rate of mice, suggesting that SteA acts as an essential effector for pathogenicity in mice.

**Fig. 4:**
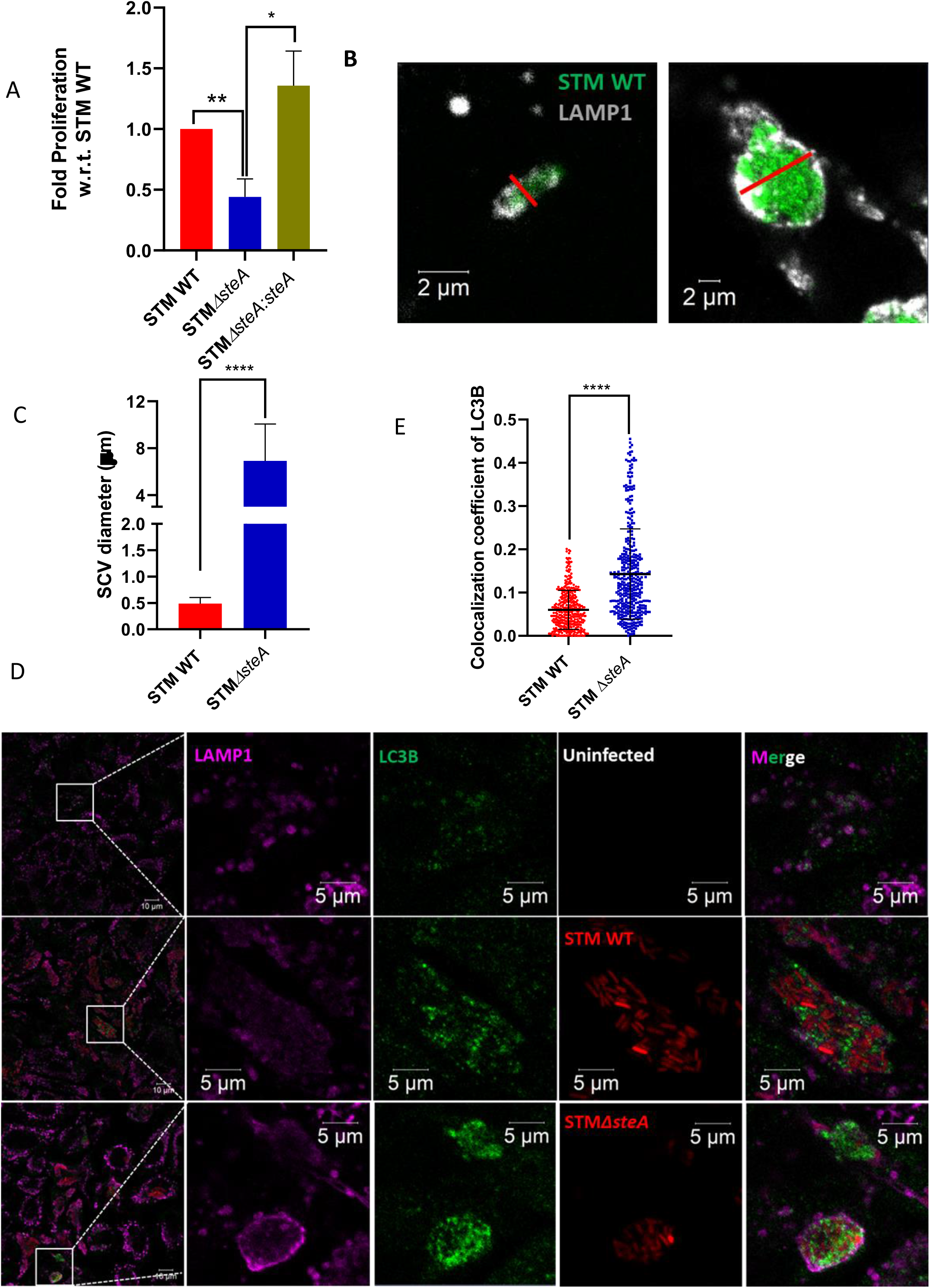
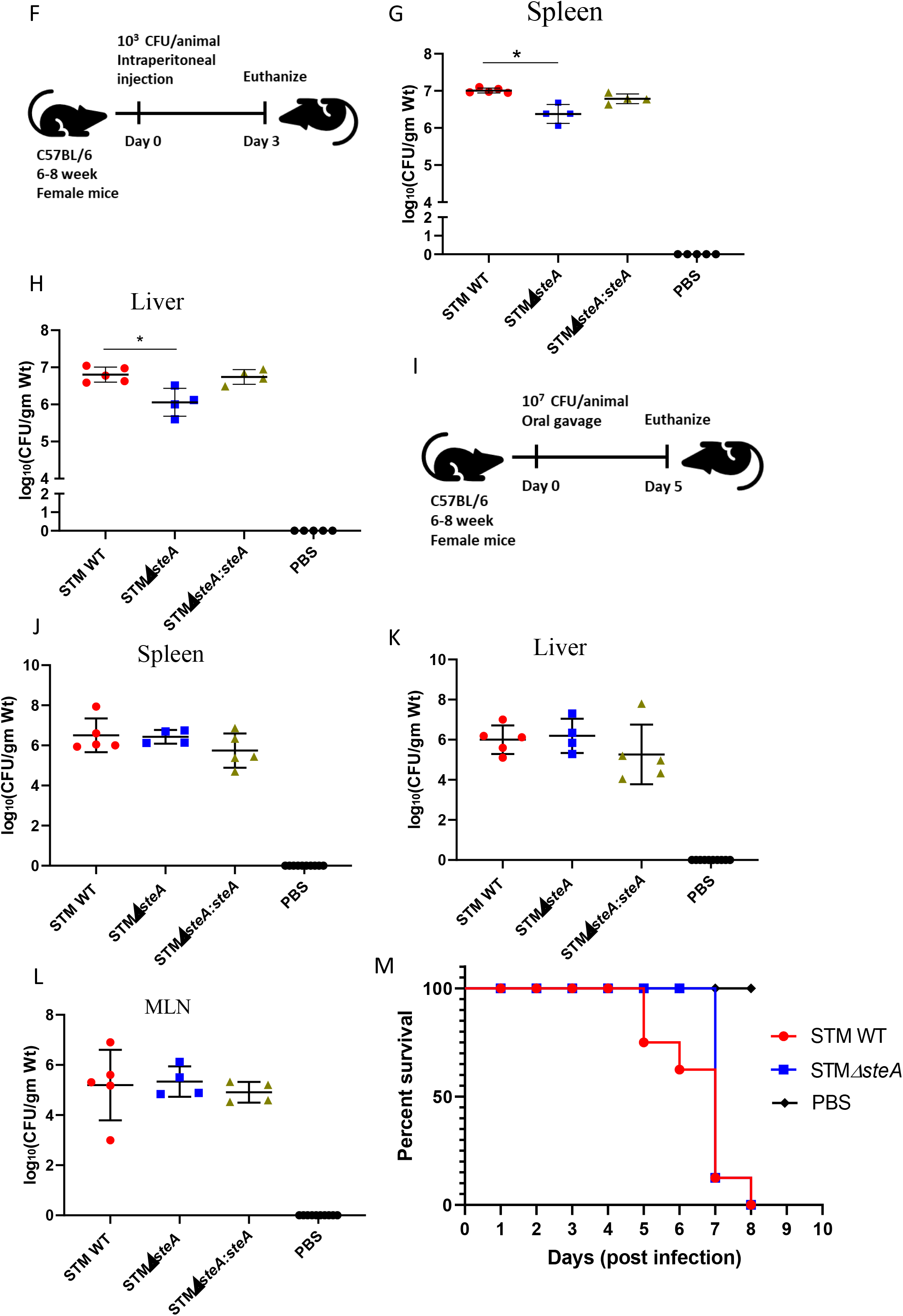
**STMΔsteA shows a defect in proliferation and pathogenicity *in vivo*** (A) Intracellular survival assay of STM WT, STM*ΔsteA*, and STM*ΔsteA:steA* in HeLa cells, data is representative of N=3,n=3, mean±SEM. (B) Line representation to quantify SCV diameter where red line indicates the length of SCV used for quantification (C) Quantification of SCV diameter, data is representative of 100-150 cells from 3 independent experiments, 16h post-infection, mean±SD (D) Representative microscopy images showing colocalization of LC3B with SCV, 16h post-infection(E) Quantification of colocalization coefficient of LC3B with SCV, data is representative of n=100-130 cells from 3 independent experiments, mean±SD (F) outline of experimental procedure of mice infection through intraperitoneal route (G) CFU enumeration in Spleen and (H) liver, data is representative of N=2, n=5 mice per cohort (I) Outline of experimental procedure of mice infection through oral gavage and dissected on day 5 post infection (J) CFU enumeration in organs like Spleen (K) Liver (L) and MLN, data is representative of N=3, n=5 mice per cohort, mean±SD (M) Survival curve of cohort of mice infected with STM WT and STM*ΔsteA,* n=8 mice per cohort. Mann-Whitney test was performed to obtain statistical significance for *in vivo* data. Student’s t-test was used to analyze the data fro m fig 4A-4E. ****<0.0001, ***<0.001, **<0.01, *<0.05.

## Discussion

Bacterial pathogens invade several cell types and reside in endocytic or phagocytic vacuoles [47]. Cell uses their extraordinary machinery to clear the pathogens, but by using several effector proteins, bacterial pathogens modulate cellular machinery and prolong their survival. A few strategies that several bacterial pathogens utilize are: modification of the membrane of the pathogen-containing vacuole, blocking the activation of xenophagy, and escaping its fusion with acidic lysosome [48], [49], [50], [51], [52], [53], [54]. Bacterial pathogens such as *Legionella, Brucella, Chlamydia*, and *Salmonella* are known to reside in vacuoles, but the morphology and membrane source of the vacuole are quite different. *Legionella* and *Chlamydia* reside in a spacious vacuole that comprises multiple bacteria in a single vacuole [55], contrary *Salmonella* resides in a tight-fitting vacuole as a single bacterium per vacuole [14]. The biogenesis of the SCV membrane is not very well known, but proteome analysis of SCV suggests that the SCV membrane is highly modified and contains proteins from the lysosome, Golgi, and ER [12]. Several effectors of *Salmonella* help in recruiting membrane components, inhibiting the transport of M6PR, and making contact sites with organelles for its proliferation [56], [57], [58]. Our results suggest that *Salmonella* infection induces ER stress inside the cell and activates the UPR. A few ways by which UPR helps in alleviating stress are 1) by increasing the protein folding capacity of cells and 2) by expanding the ER. Our data suggests that wild-type STM infection leads to UPR activation while STM*ΔssaV* is unable to activate the UPR arm. This suggests that bacterial effectors are crucial for initiating ER stress, as it was evident as early as 2h post-infection. Our studies support the previous findings, which suggest *Salmonella-*induced ER stress is crucial for bacterial proliferation [59], [60], [61]. Further, UPR activation leads to an increase in phospholipid production [40], which could be useful for the bacteria as growing SCV requires a continuous source of membrane. *Salmonella-*infected cells show an increase in expression of *Reticulon-4a* and *Climp63* with an increase in tubular ER, suggesting this is an active strategy employed by bacteria for its survival. Exogenously overexpression of Rtn4a provides an advantage in the proliferation of *Salmonella*; however, reduction in ER tubules by knockdown of Rtn4a or overexpression of ER sheets using Climp63 negatively affects the bacterial proliferation and SCV number inside the cell, indicating ER tubules plays a crucial role in bacterial proliferation. The intricate balance of ER sheets and tubules has been recently shown to affect the dynamics of membrane-less organelle [42]. Our study also highlights the importance of ER in the successful proliferation of *Salmonella*. Our live cell data showed the recruitment of ER tubules at the center of dividing SCV. This was evident in HeLa as well as RAW 264.7 macrophage cell lines, suggesting that ER tubule recruitment marks the division site of SCV and leads to a single bacterium per vacuole. ER is very well known for marking the division site of organelles such as mitochondria, membrane-less organelle, and endosomes [18], [42]. Our study suggests that *Salmonella* hijacks the strategy of ER recruitment for its survival and SCV division. We have identified the role of bacterial effector SteA in maintaining the contact of SCV with ER in the formation of possible membrane contact sites.

Earlier studies have shown the presence of SteA on SCV and its colocalization with Golgi [62], [63], [64]. In our study, we have partially observed a similar phenotype and have observed a new role of SteA in interaction with ER. Inside the infected cell, the expression of SteA is highly regulated as it can be mostly seen around the bacteria onto SCV. To investigate the role of SteA in making contact sites, we looked for the colocalization of ER with SCV of STM WT and STM*ΔsteA*. We observed the difference in colocalization of ER with SCV, highlighting the importance of *the Salmonella* effector SteA in mediating the contact site between SCV and ER. Bacterial pathogens such as *Chlamydia* use its effector protein for forming contacts of its inclusion bodies with ER [25] and it also targets host cellular protein to facilitate this interaction [65]. *Salmonella* effectors SseJ and SseL have previously been reported to recruit OSBP1 onto SCV, which helps in making contact with VapA and VapB (conserved ER proteins known for tethering of ER with cellular organelles) and maintain SCV vacuolar integrity [66]. Our data suggests the potential role of SteA in forming contact sites of SCV with ER as a strategy to facilitate its proliferation.

The loss of function of SteA led to a bulky vacuole carrying multiple bacteria inside and with defects in proliferation. Evidence from McQuate *et al*., also suggests the defect in the proliferation of STM*ΔsteA* in epithelial cells as well as macrophages [62]. To understand the difference in proliferation of STM*ΔsteA* compared to STM WT, we have observed SCV formed by STM*ΔsteA* colocalizes more with autophagy marker LC3B, suggesting big bulky vacuoles are cleared by the cellular system and hence lead to less proliferation inside the cell. STM WT also actively reduces lysosome number inside the cell as an active strategy to block its fusion with the acidic lysosome [14]. But in the case of STM*ΔsteA* infected cell, we observed a higher mean intensity of LAMP1, which was similar to the intensity observed in uninfected cells, suggesting that somehow STM*ΔsteA* could not modulate lysosomal dynamics and, as a result, active lysosome could hinder the bacterial proliferation in cell line model. Similarly, Liu *et al*.,2021 have highlighted the importance of SteA in bacterial proliferation inside cells by impairing the fusion of SCV with lysosomes [67]. Taken together, this data suggests the importance of SteA in bacterial proliferation inside infected cells. STM*ΔsteA* also showed defects in colonization in the spleen and liver if infected through the intraperitoneal route, suggesting SteA is crucial for *in vivo* pathogenicity. However, to our surprise, we did not observe any defect in the *in vivo* colonization of mouse organs such as the liver, spleen, MLN, and blood if infected through the oral gavage route. Survival analysis of mice suggests late onset of mice death in STM*ΔsteA* infected cohort upon infection as compared to STM WT. We have also observed less weight reduction in STM*ΔsteA* infected mice cohort as compared to STM WT, and this difference in weight reduction could provide an advantage in the initial survival rate of STM*ΔsteA* infected mice. (figS4D). Alongside, Geddes *et al*., 2005 have reported the importance of SteA for efficient colonization in mouse spleen using the Balb/c mice model infecting via the intraperitoneal route [63]. We have also observed similar defects in the spleen using C57BL/6 mice. However, upon infection through oral gavage, there are no defects in the colonization of STM*ΔsteA* in organs. This observation suggests that the route of infection can severely affect bacterial colonization and pathogenicity. Infection through the intraperitoneal route bypasses the gut-intestinal barrier, while during oral gavage, *Salmonella* faces several challenges, from the acidic pH of the stomach to several metabolites in the intestinal lumen. More studies are needed to understand this difference in colonization rate depending on the route of infection. Alongside, further investigations are needed to understand the vacuolar status of SCV during *in vivo* infection if infected through oral or intraperitoneal route to understand the difference in pathogenicity. Taken together, our study suggests the importance of SteA in bacterial pathogenicity and proliferation.

Our study provides new insights into how bacteria use a strategy of remodeling the ER dynamics for its benefit. Domingues et al.,2014 and Chen et al.,2021 have highlighted the importance of SteA for the active division of SCV [46], [68]. Our data supports their findings and suggests the possible role of SteA in recruiting the ER and facilitating SCV fission. Interestingly, where wild-type STM can easily divide along with SCV, while loss of function of SteA hampers 30% SCV division with marked less colocalization with ER, suggesting there can be several other effector proteins that might facilitate the contacts between ER and SCV. As reported by Chen et al.,2021 about the coordinated role of SteA, SopD2, and SifA in successful SCV division, our data explains the newly identified role of SteA in the process by making contact sites with ER. Since these effectors have coordinated roles, it will be interesting to understand how individual bacterial effectors are crucial in the complete fission of SCV. Several host proteins have also been linked with SCV division as the absence of those host proteins, such as PLEKHM1 and ESCRT-3, leads to SCV carrying multiple bacteria [57], [69]. Within the cell, SCV acts as an autonomous organelle that has acquired a vacuolar membrane from the host system, which also synchronously divides along with bacterial division. The machinery regulating the SCV division could be difficult to identify because these fission events are very transient in nature. As per literature evidence, both host and bacterial effectors could be equally involved in this process to execute successful fission. A complete knowledge of fission machinery can be used as a targeted approach to inhibit bacterial proliferation during infection.

## Acknowledgements

We acknowledge the Departmental and Divisional Confocal Microscopy Facility and Central Animal Facility at IISc for providing us with an opportunity for experimentation. Mr. Sumitlal K and Ms. Navya have helped in image acquisition in Confocal Microscopy.

## Author Contributions

UC and DC have conceived the study and designed the experiments. UC has performed all the experiments and participated in acquiring the data. UC and PB has analyzed the data, constructed the figures, and wrote the original draft of the manuscript. UC. PB, DC and GSR have participated in the proofreading and editing of the manuscript. DC has supervised the study and helped in fund acquisition. All the authors have read and approved the manuscript.

