## Supplementary figures and table for "Endoplasmic Reticulum contact sites facilitate the coordinated division of *Salmonella-*containing vacuole"

**Legends to Supplementary figure**

**Fig-S1: *Salmonella* infection leads to the activation of UPR and expansion of the Endoplasmic Reticulum**

Representative confocal microscopic images of uninfected, STM WT mCh and STM*ΔssaV* mCh infected cells, immuno-stained with (A) ATF4, LAMP1, and DAPI, 6h post-infection (B) ATF4, LAMP1, and DAPI, 2h post-infection (C) ATF6, LAMP1 and DAPI, 6h post-infection (D) ATF6, LAMP1 and DAPI, 2h post-infection (E) XBP1, LAMP1 and DAPI, 6h post-infection (F) XBP1, LAMP1 and DAPI, 2h post-infection (G) Representing uninfected, infected and bystander cell, showing an expansion of ER tubules, 16h post-infection (G) Quantification of ER tubules, data is representative of n=20-40 cells from 2 independent experiments. Student's t-test was used to analyze the data ****<0.0001, ***<0.001, **<0.01, *<0.05.

**Fig-S2: Expansion of ER tubules facilitates SCV proliferation and its division**

(A) Percent invasion of STM WT in HeLa cells upon overexpressing mCh-EV, Sec61β-mCh, and Rtn4a-mCh (B) Quantitative PCR of *Rtn4a* upon knockdown, Data is representative of N=3, n=3, mean±SD (C) Live cell imaging snapshots of RAW 264.7 murine macrophage cell line infected with STM WT showing ER overlap at fission site, N=1. Student's t-test was used to analyze the data ****<0.0001, ***<0.001, **<0.01, *<0.05.

**Fig-S3: *Salmonella* translocated effector SteA is crucial for maintaining SCV contacts with ER, resulting in a single bacterium per vacuole**

(A) Representative microscopic images showing colocalization of eGFP or SteA-eGFP with RFP KDEL (ER marker) (B) Quantification of colocalization coefficient of eGFP or SteA-eGFP with RFP KDEL. Data is representative of n=80-100 cells from 2 independent experiments, mean±SD. Student's t-test was used to analyze the data ****<0.0001, ***<0.001, **<0.01, *<0.05.

**Fig-S4: STM*ΔsteA* shows a defect in proliferation and pathogenicity *in vivo***

(A) Representative images of uninfected and infected HeLa cells with STM WT and STM*ΔsteA* showing intensity of LAMP1 (B) Quantification of mean fluorescence intensity of LAMP1, data is representative of n=100-150 cells from 3 independent experiments, mean±SD (C) CFU analysis of STM WT and STM*ΔsteA* in the blood of C57BL/6 mice, 5$^{th}$ day post oral gavage, data is representative of N=3, n=5 mice per cohort, mean±SD.(D) Represents weight reduction in mice upon infection with STM WT and STM*ΔsteA* till 8 days post-infection, 8 mice per cohort. Mann-Whitney test was performed to calculate significance. Student's t-test was used to analyze the data in fig S4B. ****<0.0001, ***<0.001, **<0.01, *<0.05.

**Supplementary File: Table: 2 List of plasmids used for overexpression.**

| Plasmid | Source |
|---|---|
| 1) pHAGE2 mCherry-Rtn4a (Plasmid-86683) | pHAGE2 mCherry-Rtn4a was a gift from Tom Rapoport (Addgene plasmid # 86683) |
| 2) mCh-Climp63 (Plasmid #136293) | mCh-Climp63 was a gift from Gia Voeltz (Addgene plasmid # 136293) |
| 3) RFP-KDEL | RFP KDEL construct was a kind gift from Prof. Nagaraj Balasubramanian |
| 4) LAMP1-GFP | GFP-LAMP1 construct was provided by Prof. Mahak Sharma |
| 5) Sec61β-mCh | mCh sec61β was a kind gift from Prof. Gia Voeltz (Addgene 49155) |

**List of shRNA plasmids used for knockdown**

| **S.No.** | **Target Gene** | **TRC ID** | **Sequence** |
|---|---|---|---|
| 1 | RTN4 reticulon 4 [ *Homo sapiens* (human) ] | TRCN0000179649 | CCGGGCAGTGTTGATGTGGGTATTTCTCGAGAAATACCCACATCAACACTGCTTTTTTG |
| 2 | RTN4 reticulon 4 [ *Homo sapiens* (human) ] | TRCN0000147624 | CCGGGCATATCTGGAATCTGAAGTTCTCGAGAACTTCAGATTCCAGATATGCTTTTTTG |

**Supplementary File: Table:3 List of Antibodies**

| Antibody | Catalog number |
|---|---|
| ATF-4 | CST-11815s |
| ATF-6 | ATF-6α Antibody (H-280): sc-22799 |
| XBP-1 | XBP-1 Antibody (M-186): sc-716 |
| LAMP1 | DSHB 1D4B |
| Anti-Salmonella | Invitrogen- PA1-20811 |
| LC3B | Sigma-L7543 |

**Supplementary File: Table-4, List of Primers used in this study:**

| **S.No** | **Primer Name** | **Sequence (5'-3')** |
|---|---|---|
| 1 | steA-KO-FP | CAAATAGTTATGGTAGCGAGCTTTTATGTCGGCCGCCCATCATATGAATATCCTCCTTAG |
| 2 | steA-KO-RP | TCAGTTTCTACCTATGCCAGAGCTTTATCAGGAAATAAGCGTGTAGGCTGGAGCTGCTTC |
| 3 | steA-eGFP cloning-FP | TTTTTCTCGAGATGCCATATACATCAGTTTCTACC |
| 4 | steA-eGFP cloning-RP | TTTTTGGATCCATAATTGTCCAAATAGTTATGG |
| 5 | HU-Rtn4a-RT-FP | TCGGGCTCAGTGGATGAGA |
| 6 | HU-Rtn4a-RT-RP | GCAGGACAGATGGGAAATCCT |
| 7 | HU-Climp63-RT-FP | GCGTCGAGCAGAAGGTGC |
| 8 | HU-Climp63-RT-RP | CATGGATCCCATCCGAGAGG |
| 9 | Xho1_fwd_Native_promoter_steA_cloning | TTTTTCTCGAGCGGCAGTGATTGCGTTGC |
| 10 | HindIII steA-HA_cloning_RP | TTTTTTAAGCTTTTATGCATAATCCGGAACATCATACGGATAATAATTGTCCAAATAGTTATGGTAGCGAGC |
| 11 | HU-β-actin-FP | GCCGCCAGCTCACCAT |
| 12 | HU-β-actin-RP | TCGTCGCCCACATAGGAATC |

**Fig S1: *Salmonella* infection leads to the activation of UPR and expansion of the Endoplasmic Reticulum**

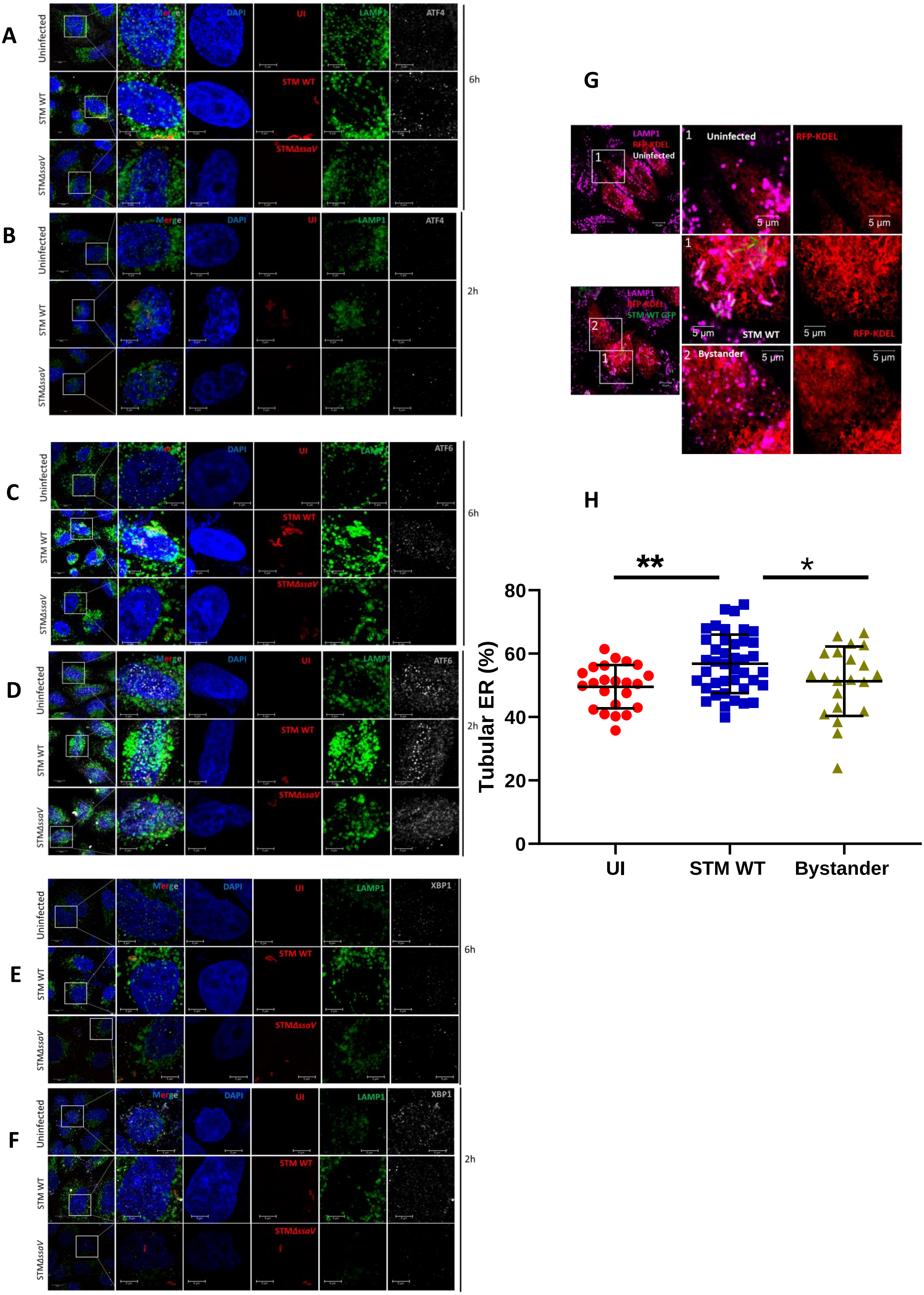

**Fig S2: Expansion of ER tubules facilitates SCV proliferation and its division**

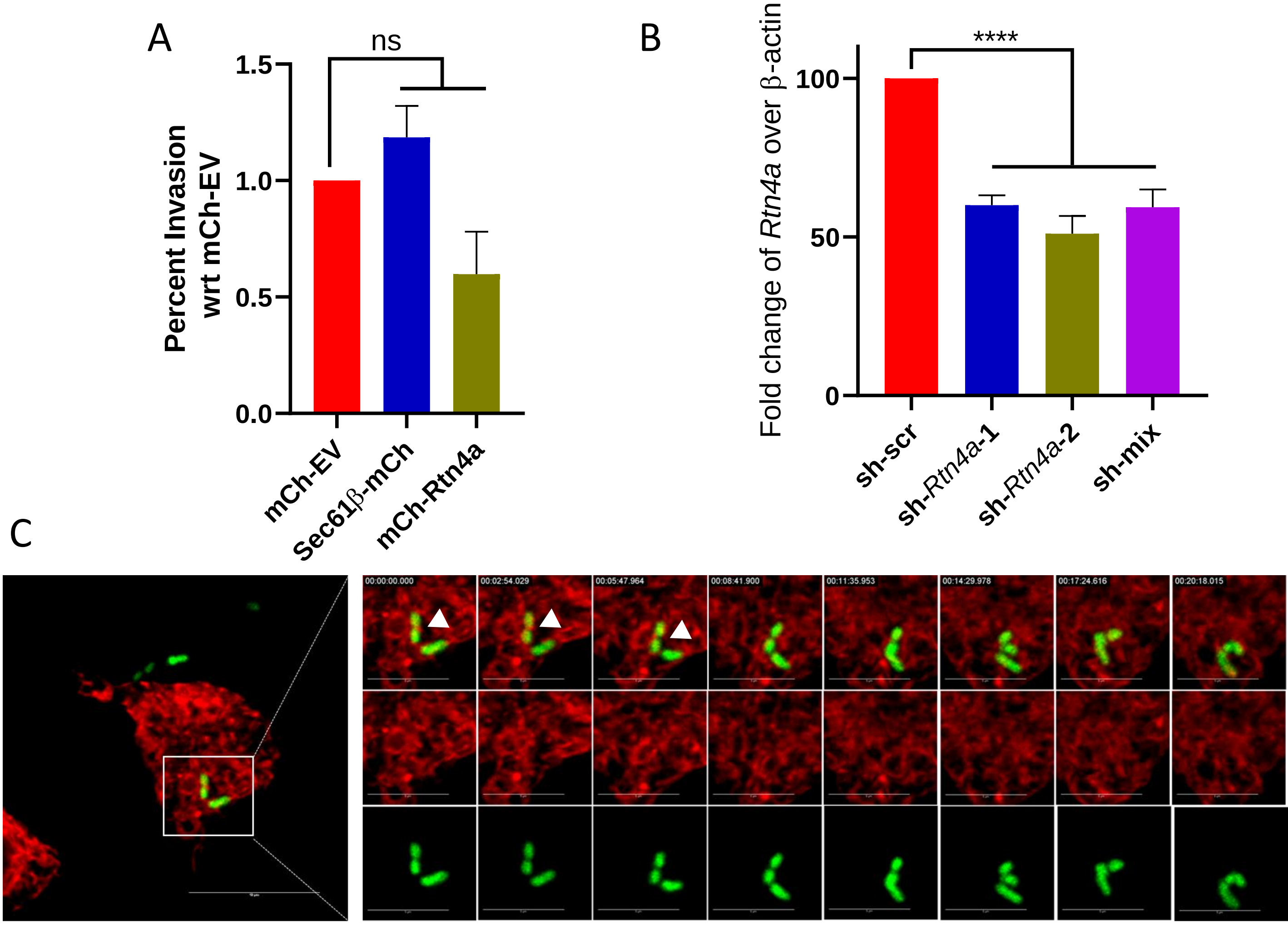

**Fig S3: *Salmonella* translocated effector steA is crucial for maintaining SCV contacts with ER resulting in a single bacterium per vacuole state**

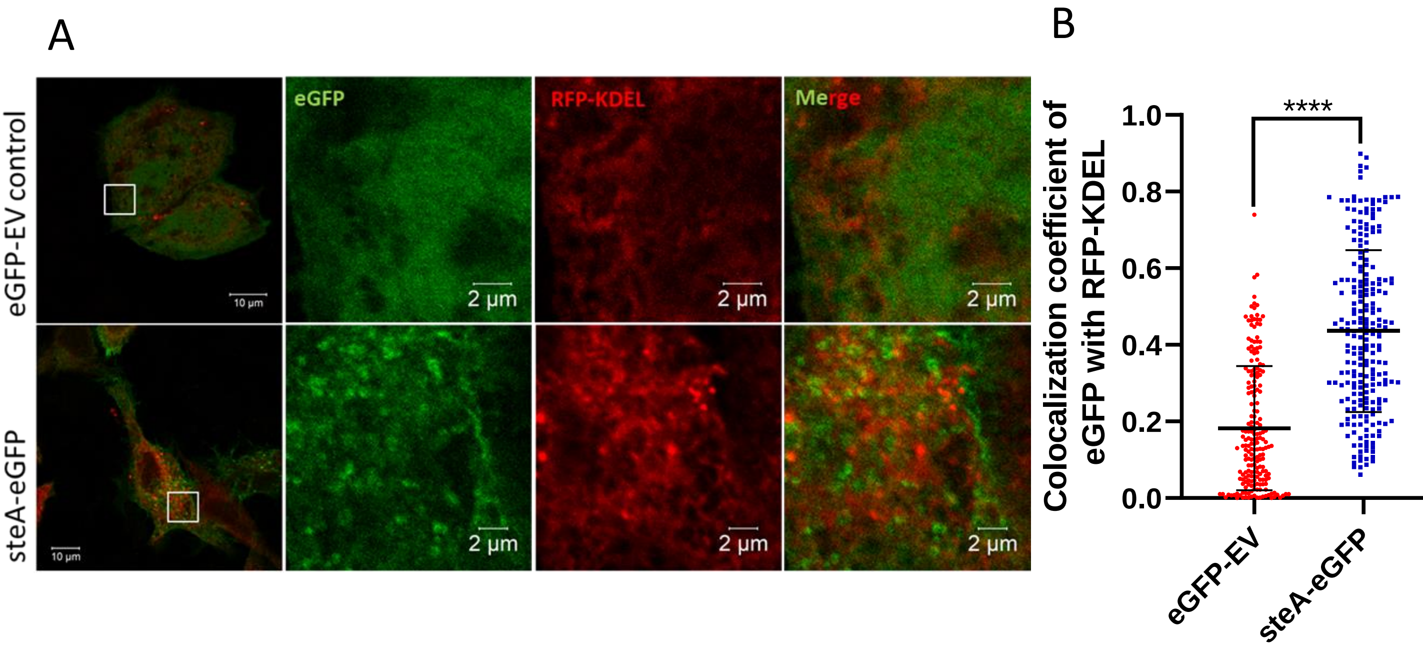

**Fig S4: STM$\Delta$*steA* shows a defect in proliferation and pathogenicity *in vivo***

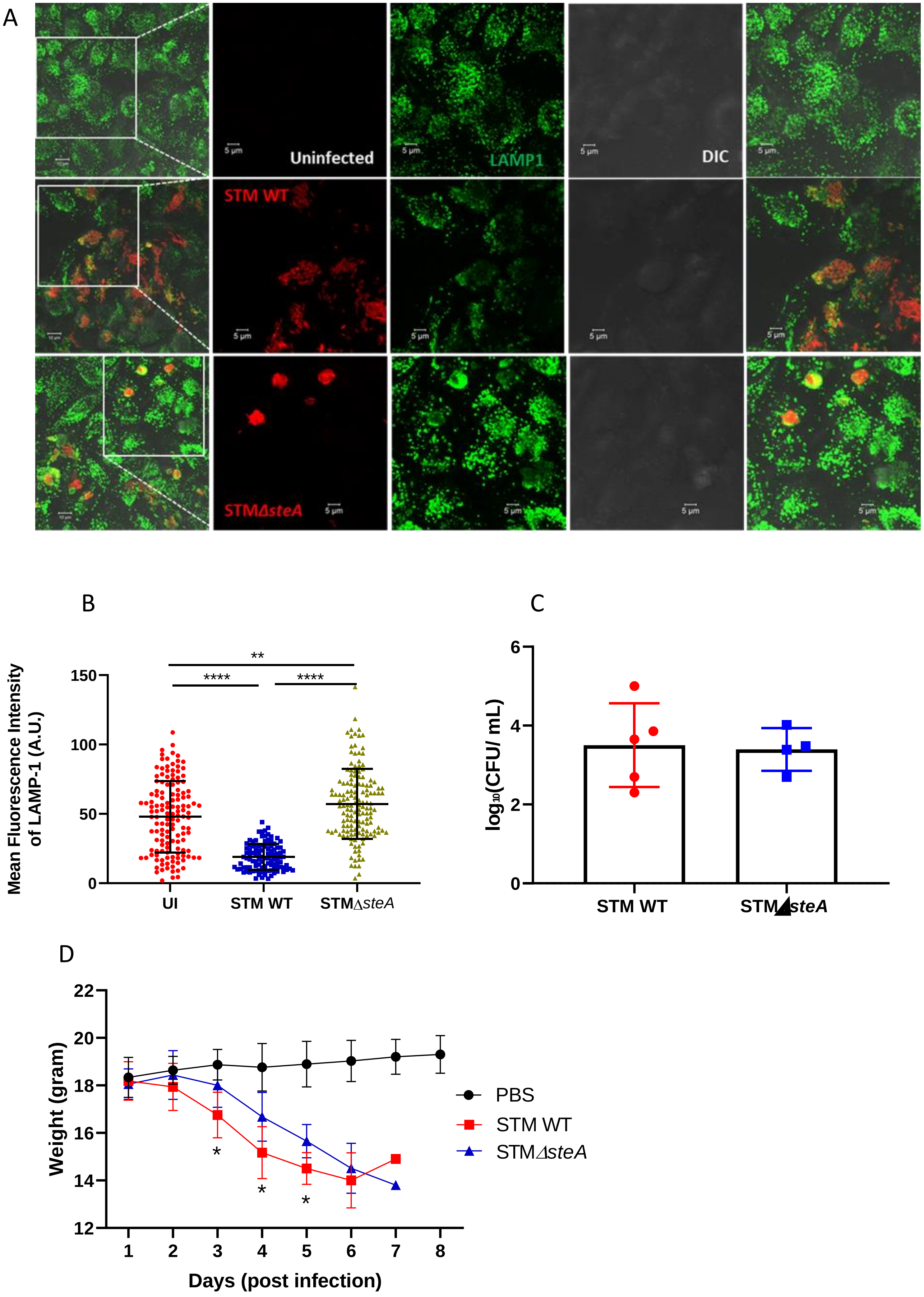
